# Integrating bulk and single cell RNA-seq refines transcriptomic profiles of individual *C. elegans* neurons

**DOI:** 10.1101/2025.01.26.634951

**Authors:** Alec Barrett, Erdem Varol, Alexis Weinreb, Seth R. Taylor, Rebecca M. McWhirter, Cyril Cros, Berta Vidal, Manasa Basaravaju, Abigail Poff, John A. Tipps, Maryam Majeed, Chen Wang, Emily A. Bayer, Molly Reilly, Eviatar Yemini, HaoSheng Sun, Oliver Hobert, David M. Miller, Marc Hammarlund

## Abstract

Neuron-specific morphology and function are fundamentally tied to differences in gene expression across the nervous system. We previously generated a single cell RNA-seq (scRNA-Seq) dataset for every anatomical neuron class in the *C. elegans* hermaphrodite. Here we present a complementary set of bulk RNA-seq samples for 52 of the 118 canonical neuron classes in *C. elegans*. We show that the bulk RNA-seq dataset captures both lowly expressed and noncoding RNAs that are not detected in the scRNA-Seq profile, but also includes false positives due to contamination by other cell types. We present an analytical strategy that integrates the two datasets, preserving both the specificity of scRNA-Seq data and the sensitivity of bulk RNA-Seq. We show that this integrated dataset enhances the sensitivity and accuracy of transcript detection and differential gene analysis. In addition, we show that the bulk RNA-Seq data set detects differentially expressed non-coding RNAs across neuron types, including multiple families of non-polyadenylated transcripts. We propose that our approach provides a new strategy for interrogating gene expression by bridging the gap between bulk and single cell methodologies for transcriptomic studies. We suggest that these datasets advance the goal of delineating the mechanisms that define morphology and connectivity in the nervous system.

## Introduction

Neurons exhibit an extraordinary range of morphological forms and physiological functions. Because this diversity is largely driven by underlying differences in gene expression, a key goal of neuroscience is to identify the transcripts expressed in each neuron type.

The adult *C. elegans* hermaphrodite contains 302 neurons divided into 118 anatomically distinct neuron types. The structure, connectivity, and lineage are known for each of these neurons(*1–6*). Recently, the *C. elegans* Neuronal Gene Expression Map & Network project (CeNGEN)(*7*) used single cell RNA sequencing (scRNA-seq) technology to generate a gene expression atlas that matches the single neuron resolution of the structural map of the mature *C. elegans* nervous system(*8*).

The CeNGEN scRNA-seq dataset was acquired with 10x Genomics technology and is largely comprised of reads from poly-adenylated transcripts. Thus, major classes of non-poly-adenylated transcripts, especially noncoding RNAs, are under-represented in the CeNGEN scRNA-seq data. In addition, low abundance transcripts may be sparsely detected in scRNA-seq data, particularly in clusters with relatively few cells(*8*). Both noncoding RNAs and low abundance transcripts are potentially important mediators of neuronal fate. A description of their expression is therefore needed to complement the CeNGEN scRNA-seq map of neuronal poly-adenylated transcripts.

Here, we use FACS to isolate single neuron types for bulk RNA sequencing with the goal of describing neuronal gene expression with high sensitivity and specificity. We generated profiles for 53 individual neuron types from the mature *C. elegans* hermaphrodite nervous system. These 53 types include 52 of the canonical 118 types, with separate data for the subclasses ASEL and ASER. This data set samples a wide range of neuron types including motor neurons, interneurons, and sensory neurons. We built sequencing libraries with random primers for robust detection of both poly-adenylated and non-coding RNAs(*9*). We developed computational approaches that exploit the bulk dataset to enhance accurate detection of gene expression, including integration with the existing CeNGEN scRNA-seq dataset and methods for measuring non-coding RNAs. The resulting data sets refine quantitative measures of gene expression and improve detection of low-abundance and non-poly-adenylated transcripts. These data provide a unique opportunity for future studies that link gene expression to neuron function, structure, and connectivity.

## Methods

### Strains

Strains used for FACS isolation of individual neuron classes are listed in Table 1. Strains were either generated by us or are kind gifts of the *C. elegans* community. The lab origin of individual transgenic arrays can be inferred from the allele designation, as detailed at Lab List at the CGC (https://cgc.umn.edu/laboratories). All strains used in this study are available at the CGC.

**Table 1:**
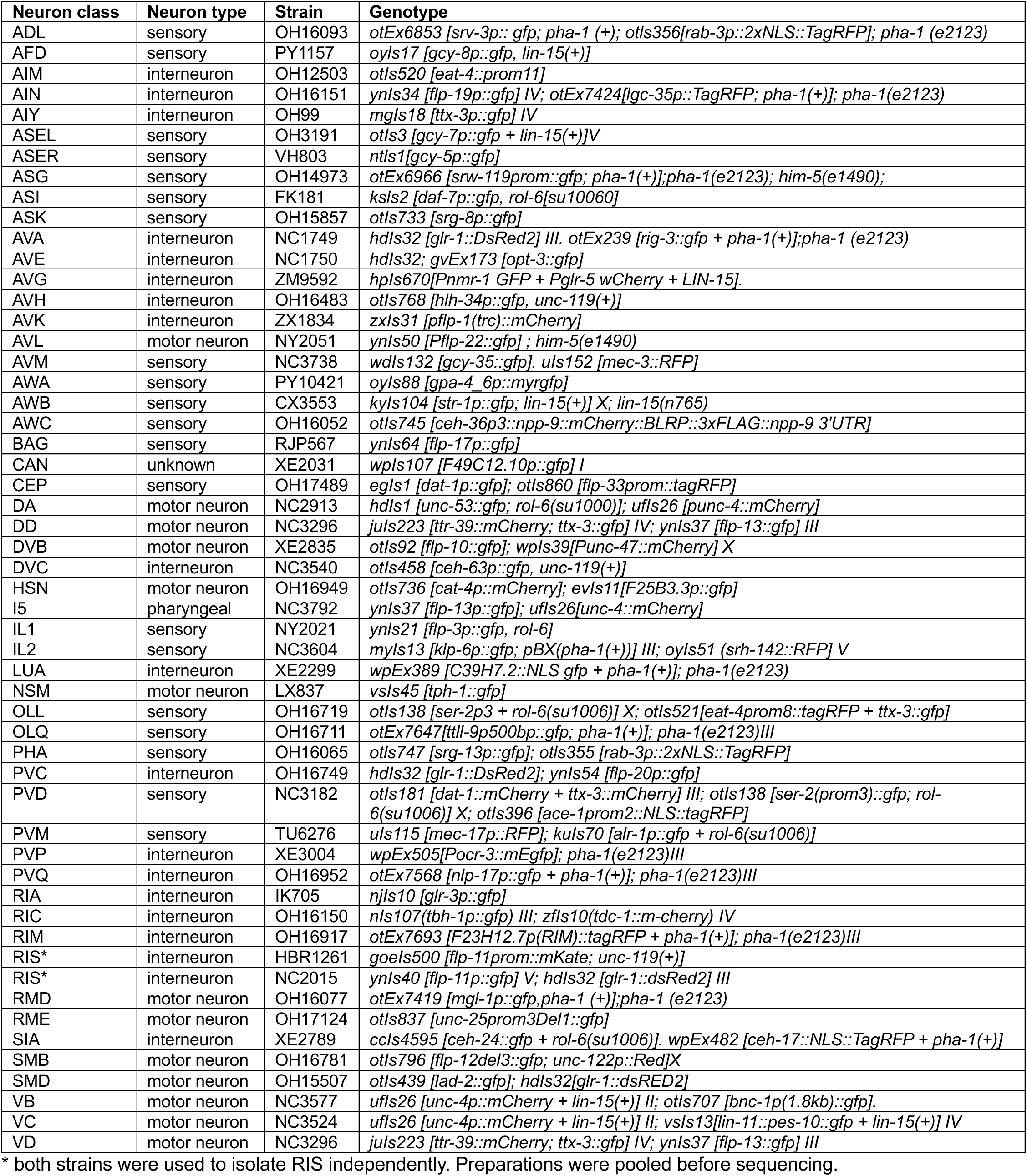
Transgenic strains used for isolating individual neuron classes.

### FACS isolation for RNA-seq

Labeled neuron types were isolated for RNA-seq as previously described(*10–12*). Briefly, synchronized populations of L4 stage larvae were dissociated and labeled neuron types isolated by Fluorescence Activated Cell Sorting (FACS) on a BD FACSAria III equipped with a 70-micron diameter nozzle. DAPI was added to the sample (final concentration of 1 mg/mL) to label dead and dying cells. For bulk RNA-sequencing of individual cell types, sorted cells were collected directly into TRIzol LS. At ∼15-minute intervals during the sort, the sort was paused, and the collection tube with TRIzol was inverted 3-4 times to ensure mixing. Cells in TRIzol LS were stored at -80C for RNA extractions. See Table S1 for list of samples and associated metadata.

### RNA extraction

RNA extractions were performed as previously described(*8, 12*). Briefly, cell suspensions in TRIzol LS (stored at -80°C) were thawed at room temperature. Chloroform extraction was performed using Phase Lock Gel-Heavy tubes (Quantabio) according to the manufacturer’s protocol. The aqueous layer from the chloroform extraction was combined with an equal volume of 100% ethanol and transferred to a Zymo-Spin IC column (Zymo Research). Columns were centrifuged for 30 s at 16,000 RCF, washed with 400 mL of Zymo RNA Prep Buffer, and centrifuged for 16,000 RCF for 30 s. Columns were washed twice with Zymo RNA Wash Buffer (700 mL, centrifuged for 30 s, followed by 400 mL, centrifuged for 2 minutes). RNA was eluted by adding 15 mL of DNase/RNase-Free water to the column filter and centrifuging for 30 s. A 2 μL aliquot was submitted for analysis using the Agilent 2100 Bioanalyzer Picochip to estimate yield and RNA integrity, and the remainder was stored at -80°C. See Table S1 for list of samples and associated metadata.

### Bulk sequencing and mapping

Each bulk RNA sample was processed for sequencing using the SoLo Ovation Ultra-Low Input RNaseq kit from Tecan Genomics according to manufacturer instruction, modified to optimize rRNA depletion for *C. elegans*(*9*). Libraries were sequenced on the Illumina Hiseq 2500 or Novaseq6000 with 150 bp paired end reads. Reads were mapped to the *C. elegans* reference genome from WormBase (version WS289) using STAR (version2.7.7a) with the option --outFilterMatchNminOverLread 0.3. Duplicate reads were removed using UMI-tools (v1.1.4), and a counts matrix was generated using the featureCounts tool of SubRead (v2.0.3). FASTQC was used for quality control before alignment, and four samples were removed for failing QC or for a low number of reads.

Each biological sample contains one or more transgenes, and each transgene generally consists of many copies of a promoter sequence driving expression of a fluorophore (Table 1). To avoid artifacts stemming from spurious transcription of high-copy number transgenes, we masked 5kb of all genes whose promoters were used in transgenes, starting 4kb upstream of the start ATG and extending 1kb past the ATG. This approach removed 231 genes from our analysis (Table S2).

### Sample Normalization

Intra-sample normalization (gene length normalization for bulk samples) was performed before integration (see below). Inter-sample normalization (library size normalization) was performed after integration. Library size normalizations were performed using a TMM (trimmed mean of M-values) correction in edgeR (version 4.0.1).

### Ground-truth genes

As an independent measure of gene expression, we used a “ground truth” dataset of 160 genes for which expression in individual neuron types is known with high precision across the entire nervous system (Table S3). This expression matrix is based on high confidence fosmid fluorescent reporters, CRISPR strains or other methods(*8, 13–17*).

We also curated a list of 445 genes that are exclusively expressed outside the nervous system to assess potential non-neuronal contamination in each sample (Table S4). This list was curated from published datasets of fluorescent reporters, tissue specific RT-PCR, and transcriptomic studies available on WormBase(*15*). Genes were included if two types of evidence documented expression in the same non-neuronal tissue (non-identical expression in other tissues was allowed so long as at least one tissue overlapped), and for which there was no evidence for neuronal expression.

An additional 936 ubiquitously expressed genes were used to evaluate gene expression accuracy in the bulk RNA-Seq samples (Table S5). To compile this list, we used the scRNA-Seq data(*11*), and defined the expression level of a gene in a given cluster as the proportion of cells in that cluster in which at least one UMI was detected. We then defined a gene as ubiquitous if at least 10 clusters displayed an expression level of ≥0.5%, excluding neuron-specific genes.

### Comparing datasets to ground-truth

When comparing data to “ground truth” gene expression, a static threshold was applied to the average normalized cell profile (i.e., arithmetic mean across all cells, or samples). Single cells were normalized to library size prior to averaging to calculate TPM counts(*18*). Bulk samples were normalized with the TMM method using edgeR (version 4.0.1), noncoding RNA analysis was performed with length normalized TMM counts using the GeTMM method(*19*).

### Subtraction

bMIND(*20*) was run by first excluding genes that are invariant in all cell types. Normalized aggregate tissue level scRNA-seq profiles for each cell type were used as the prior reference. Cell-type proportions were estimated using the NNLS option.

ENIGMA(*21*) was run by first excluding genes that are invariant in all cell types. Normalized aggregate tissue-replicate level scRNA-seq profiles for each cell-type were used as the reference, meaning that each tissue was represented in multiple columns, with different columns for each experimental replicate in the scRNA-seq dataset. Cell-type proportions were estimated using the robust linear regression (RLR) option.

ENIGMA was run using the L2 norm, with a log transformation preprocessing step. Trace norm runs did not improve performance on ground truth gene detection (data not shown).

LittleBites (this work) was run by first calculating gene level specificity weights in a single cell reference dataset using the SPM method(*22, 23*). The SPM method weights genes highly (values close to 1) only if they are present exclusively in one cell type, and the method is robust to different numbers of cell types. SPM specificity scores are conservative, and result in less subtraction for any gene that is expressed even in just a few cell types. The LittleBites algorithm runs in 4 steps: 1) Model each bulk sample as a linear combination of single cell reference profiles of target and putative contaminant tissues, using a non-negative least squares approach. The estimated coefficients are used as estimates of the bulk sample composition. 2) Subtract the non-target single cell reference profiles from the bulk sample, using the modeled coefficients as weights for each corresponding profile (Equation 1). For each subtraction, set a range of learning rates from 0 to 1, and perform the subtraction once for each learning rate supplied. 3) For each learning rate, compare the subtracted values against known ground truth genes, and calculate the AUROC, and return the subtracted sample that maximizes the AUROC value. If more than one learning rate returns the same AUROC value, return the subtracted dataset with the lowest learning rate. 4) If there is no learning rate that produces a higher AUROC than a learning rate of 0, halt subtraction. Else, return to step 1 using the subtracted bulk sample as the input.

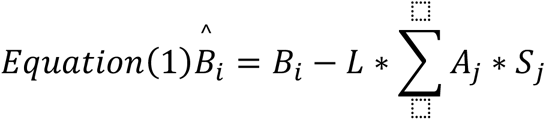

B = bulk sample, A = estimated composition coefficient, S = Single cell reference, L = learning rate.

Cell-type level aggregate data from the scRNA-seq dataset were used as a reference, with just the neuron of interest used to model each sample. LittleBites was run with a log transformation preprocessing step for bulk and single cell references. LittleBites code is available at https://github.com/alecbarrett/LittleBites.

### Bulk sample composition estimates

Contamination estimates were performed for each bulk sample by using non-negative least squares (NNLS) modeling on down-sampled and log transformed counts, averaging across 100 estimates per sample. Down-sampling was performed to reduce bias against neuron types with small cluster sizes. For each bulk sample (ex: AFD replicate 1), proportions were estimated using only neuronal cells for the corresponding single cell cluster (ex: AFD), and 7 identified non-neuronal clusters (Glia, Excretory, Hypodermis, Intestine, Muscle-mesoderm, Pharynx, and Reproductive). For each iteration, all 8 single cell clusters were down-sampled to 30 cells each, and average TPM counts were calculated using the arithmetic mean for each gene in the 30 cells.

Gene level variance was calculated using the averaged TPM values, and low variance genes were removed. Bulk sample counts and single cell TPMs were log transformed before the NNLS calculation with a pseudocount of one. NNLS estimates across all 100 iterations were averaged for the final estimate. NNLS calculations were performed using the nnls package in R (version 1.5).

### Correlating gene expression to non-neuronal contaminants

Each gene was correlated to non-neuronal contamination across all samples using Spearman’s correlation test. High correlation to any contaminant was used to indicate that the gene is likely detected because of contamination, and not due to expression in the target neuron. For genes passing an expression threshold > 2 normalized counts in at least 2 samples, their highest correlation value to any contaminant tissue was collected, and cutoffs were determined by fitting a gaussian mixture model using the normalmixEM2comp function in mixtools (version 2.0.0), fitting 2 gaussian distributions to the distribution of highest contaminant correlations. Cutoffs were selected to exclude 95% of the predicted contaminant distribution.

### Defining new gene expression for lowly expressed genes and for noncoding genes

For protein coding genes that are under the detection threshold for specific single cell clusters, but are detectable elsewhere in the single cell dataset, we identified expanded gene expression profiles using the LittleBites-cleaned bulk dataset. These genes were thresholded (>40.3 TMM) to match the FDR (14%) for the published single cell analysis(*8*). For protein coding genes that are never detected in the single cell dataset, a threshold was set using the FPR for ground truth non-neuronal genes, after excluding genes with high non-neuronal contaminant correlation values. These genes were called expressed if they were detected above 6.06 TMM, which corresponds to an FPR of 0%. For noncoding RNAs, genes with high correlation to non-neuronal contaminant estimates were removed, and for the remaining genes the threshold was set at 5 normalized counts, and genes were called expressed if they passed that threshold in >65% of samples.

### Pseudobulk aggregation of single-cell data

We downloaded the CeNGEN scRNA-seq dataset as a Seurat object from the CeNGEN website (www.cengen.org). Cells from the same cell type and experiment (e.g. AFD cluster) were aggregated together by summation into a single pseudobulk sample. For this work, single cell clusters of neuron subtypes were collapsed to the resolution of the bulk replicates (example: VB and VB1 clusters in the single cell data were treated as one VB cluster). Pseudobulk samples from cluster-replicates with fewer than 10 cells were excluded.

### Modeling Single Cell cluster proportions and counts

To simulate a single cell pseudobulk-cluster, we sampled from a negative binomial distribution for each gene using the rnbinom function from the stats package in R (Version 4.3.2). We then modeled the number of counts per gene as the sum of the log odds-ratio and the log of the total cells in the simulated cluster (equation 2). For a given cluster k and gene g, C_k,g_ = total counts for the gene in the cluster, P_k,g_ = proportion of cells with at least 1 count for a given gene, and n_k_ = the total cells in the cluster

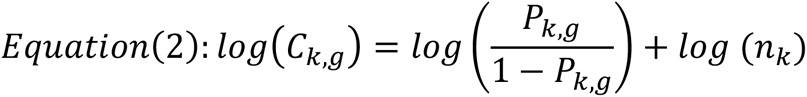

We applied this formula to our real single cell dataset and used this equation to transform proportion measures of gene expression into a count space to generate the Prop2Count dataset for downstream analysis and integration with bulk datasets. This procedure allows for proportions data to be used in downstream analyses that work with counts datasets. However, it does limit the range of potential values that each gene can have, with the potential values set as:

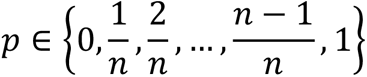

As n approaches 0, the number of potential values decreases, which can be incompatible with some downstream models. Thus, caution should be used when applying this transformation to datasets with few cells.

Prop2Count code is available at https://github.com/alecbarrett/Prop2Count.

## Results

### Bulk sequencing of individual neuron types

The model organism *C. elegans* is uniquely suitable for the task of defining gene expression in the nervous system at high resolution and genome scale. *C. elegans* is the first metazoan with a completely sequenced genome(*24*) and the only animal for which we know every cell division that gives rise to the adult body plan (*i.e.,* “cell lineage”)(*1, 2*), as well as the anatomy of each neuron and all of its connections with other cells(*3–6, 25*). The entire *C. elegans* hermaphrodite nervous system contains 302 neurons with 118 canonical anatomically-defined neuron classes, each comprised of relatively few cells, ranging from 1 to 13 neurons(*3*). Most of these neuron classes are either a bilateral pair of anatomically similar cells (70 classes) or single neurons (26 classes) with unique morphological and functional characteristics. The rich array of distinct neuron classes in *C. elegans,* combined with the fact that these types are invariant among individuals, means that each neuron class can be analyzed in depth to reveal the genetic programs that define neuronal diversity.

We previously generated a gene expression atlas for the entire *C. elegans* nervous system at the resolution of single neuron types(*8*). We completed this atlas with single-cell techniques by adopting the strategy of using FACS to enrich for specific groups of neurons for a series of scRNA-seq experiments. The single-cell atlas provides a detailed description of gene expression across the *C. elegans* nervous system.

However, the single-cell atlas may fail to detect lowly-expressed genes, particularly in clusters with few cells. Also, due to its reliance on poly-dT priming for reverse transcription, the single-cell atlas excludes non-coding transcripts that are not poly adenylated(*11*) (see Figure 5B). Finally, because the scRNA-seq data is biased toward the 3’ ends of transcripts, it does not contain information about most splicing events(*26, 27*).

To address these limitations and to provide a broader description of gene expression across the nervous system, we developed a bulk RNA sequencing strategy to profile individual neuron types(*10*). We hypothesized that bulk RNA-seq and scRNA-seq datasets might have complementary strengths and weaknesses. Bulk RNA-seq can enhance sequencing depth and gene detection, capture non-polyadenylated transcripts, and provide uniform coverage of the transcript body(*9*). However, bulk RNA-seq data are typically contaminated with transcripts from non-target cell types which can limit specificity for some genes. By contrast, scRNA-seq datasets allows for high specificity in gene detection, as contaminating cells can be identified post-hoc, but can show reduced transcript sensitivity, especially for low abundance cell types(*8*).

Our bulk sequencing strategy uses a series of *C. elegans* strains, each of which contains one or more fluorescent markers that label an individual neuron type for isolation by FACS. For example, we used *flp-22*::GFP and *unc-47*::mCherry, which together uniquely mark mark the single neuron AVL, for FACS (Fig. 1A). We selected strains based on the following criteria: 1) In the desired neuron type, the fluorophore(s) is expressed brightly enough to be clearly visible under a stereo scope. 2) The fluorophore, or combination of fluorophores, is uniquely expressed in the desired neuron type. These strains thus represent a rich resource for studying individual neuron types in *C.* elegans; all are deposited at the CGC (Table 1).

**Figure 1:**
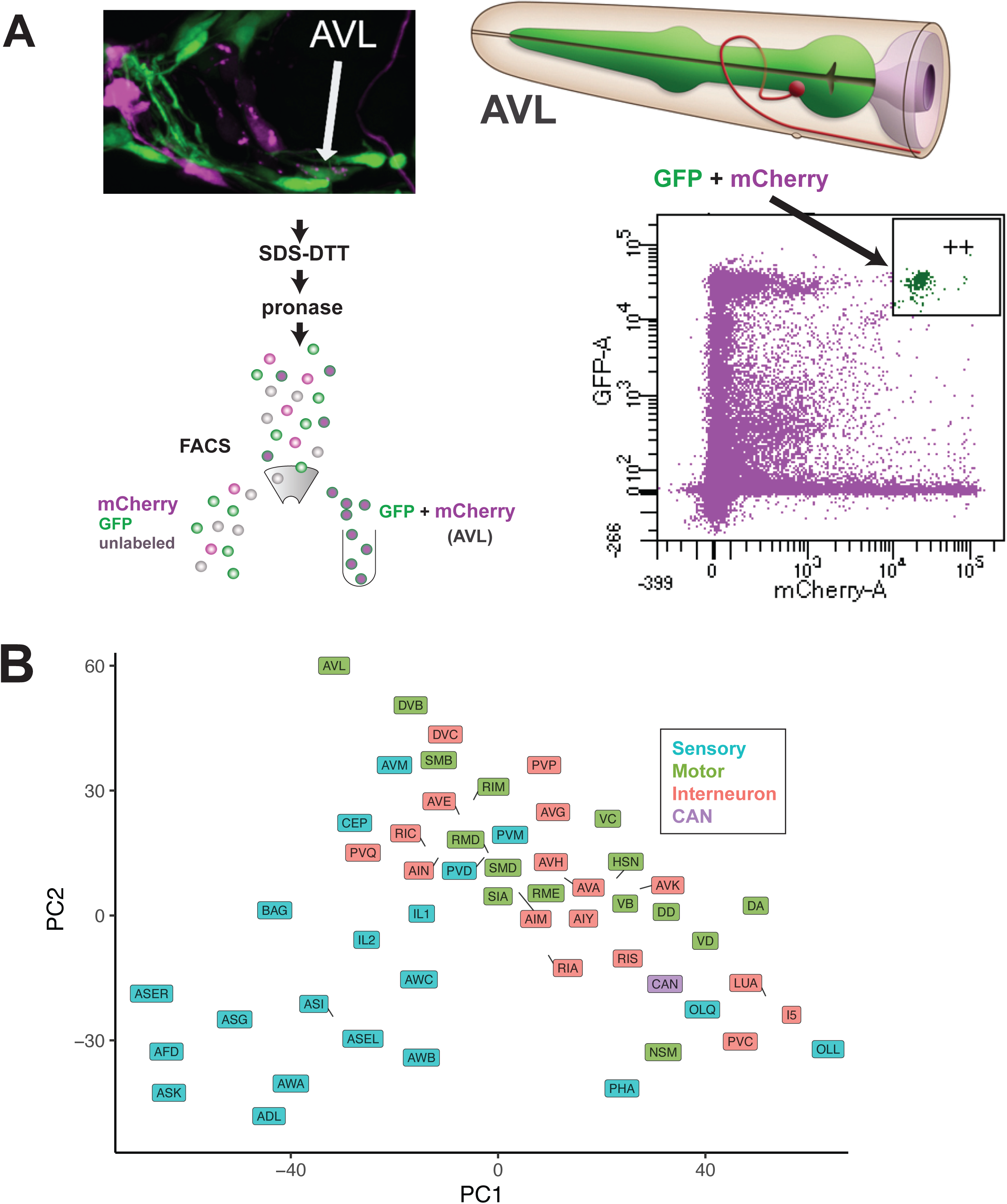
Single neuron bulk RNA-seq via targeted marker expression and FACS isolation. A) Labeling, tissue dissociation, and FACS-enrichment schemes for capturing individual neuron types. Intersecting *flp-22*::GFP and *unc-47*::mCherry markers uniquely label AVL for isolation by FACS from dissociated L4 stage larval cells. RNA from this pool of AVL-enriched cells was used for bulk RNA sequencing (see Methods). B) PCA plot showing all bulk RNA-seq data labeled by cell type and colored according to functional modality; Sensory neurons (blue), motor neurons (green), interneurons (red), and CAN neurons (purple).

For each strain, we used FACS to isolate neurons from synchronized populations of hermaphrodites at the L4 stage, by which time all neurons have been born and are differentiated. Labeled cells were collected in TRIzol LS for RNA extraction. We isolated a wide range of cells (∼700 – 90,000) in each sample across neuron types. Multiple biological replicates (e.g., separately grown cultures that were then dissociated and sorted separately) were generated for each neuron class. In total, we sequenced 206 samples across 53 neuron types (Table S1). The 53 neurons that we profiled sample a wide range of anatomical locations (head ganglia, ventral cord, mid-body and tail neurons, pharyngeal neurons) functional modalities (sensory, inter- and motor neurons), neurotransmitter usage (glutamatergic, GABAergic, cholinergic, aminergic) and lineage history (Fig. S1A). A few of these bulk neuron profiles have been previously described(*11*).

We used a ribodepletion strategy combined with random priming for cDNA synthesis. This approach optimized whole transcript coverage for each gene and also captured non-polyadenylated RNAs (see Methods)(*9*). The resultant datasets comprise a high-resolution view of RNA expression across the *C. elegans* nervous system. A distribution of neuron-specific data sets for the first two principal components shows separation between sensory neurons (especially ciliated sensory neurons, e.g. ASK, ADL, AFD, AWA) vs motor and interneurons, a result consistent with patterns observed for scRNA-seq data on the same neuron classes (Fig. 1B)(*8*). The average bulk data for each neuron type also correlates with the single-cell data for that neuron type, with the exception of the OLL neuron, which is an outlier in the single cell dataset with few sampled cells, and may therefore be less accurate (Fig. S1B). Overall, these new bulk datasets enable new studies of neuronal gene expression in individual neuron types in *C. elegans*.

### A novel subtraction approach to clean bulk data using information from single-cell data

Our bulk datasets rely on FACS for enrichment of each specific neuron type. Although FACS-isolated cell populations are highly enriched for the target cell type, these samples are also typically contaminated with additional RNA from other cell types— either from errant cells or from ambient RNA in the cell suspension(*28, 29*). To address this problem, we developed an approach, ‘LittleBites’, that uses pre-existing and reliable information on gene expression to remove contamination from bulk sample data.

LittleBites performs iterative linear subtraction on bulk RNA-Seq data using an scRNA-seq reference and a set of ground truth genes. In brief, LittleBites models each bulk sample as a mix of single cell expression profiles, estimates the cell-type proportions, and then subtracts the non-target profiles. The subtraction is compared to a set of ground truth genes, and repeated iteratively until no further improvements can be made (Fig. 2A; see Methods). For example, the bulk VB sample VBr31 had an initial composition estimate of: 22% neuronal, 0% excretory, 8% glia, 13% hypodermis, 17% intestine, 4% pharynx, 0% muscle, and 35% reproductive, and its area under the Receiver-Operator Characteristic (AUROC) score for ubiquitous and non-neuronal testing genes was 0.86. After LittleBites subtraction, the composition estimates were: 88% neuronal, 2% excretory, <1% glia, 0% hypodermis, 0% intestine, <1% pharynx, 6% muscle, and 2% reproductive, and the AUROC score for the testing gene set increased to 0.96. On average, the neuronal estimate for each sample increased 2.8-fold, and the testing gene AUROC increased by 0.084 (9.5%).

**Figure 2:**
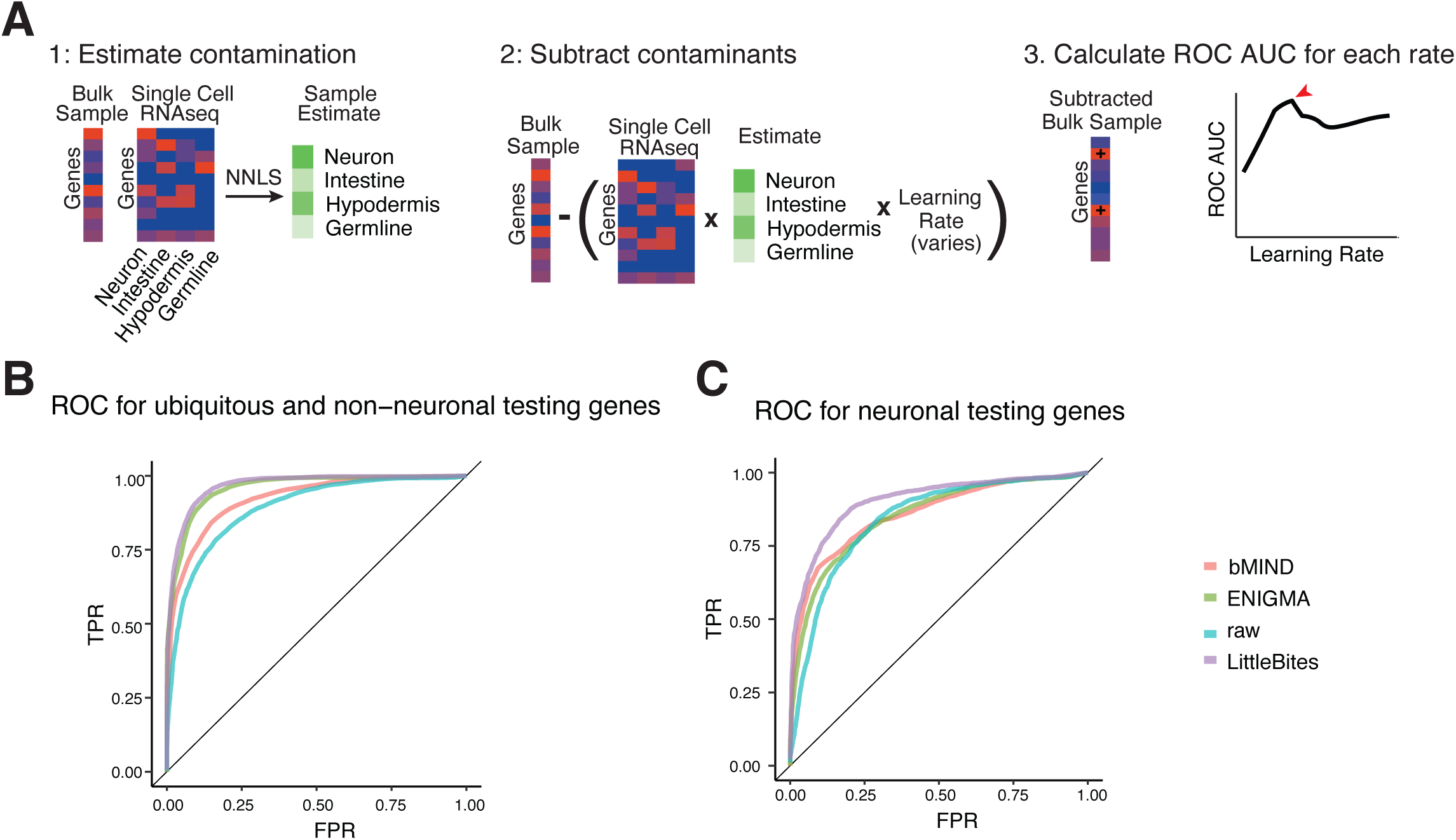
Identifying and removing contamination from bulk data with LittleBites. A) Schematic representation of the LittleBites algorithm. Each iteration of the LittleBites algorithm contains 3 steps: 1) Use an NNLS regression to estimate the cell-type proportions of each bulk sample with aggregated single cell data as a reference. 2) Subtract the estimated contamination profiles, constrained by the proportion estimates, a gene level specificity weight (not shown), and a variable learning rate. 3) Compare the result of every tested learning rate against ground truth, and select the learning rate with the highest area under the receiver operator characteristic (AUROC). This loop repeats until further subtraction cannot improve the AUROC. B) Line plots of the receiver operator characteristic (ROC) curve for the unaltered bulk data and corrected datasets, as well as bMIND and ENIGMA analyses, assessed against ubiquitous genes and genes exclusively expressed in non-neuronal cells. C) Line plots of the ROC for the unaltered bulk data and corrected datasets, as well as bMIND and ENIGMA analyses, assessed against 160 genes expressed within the nervous system.

To assess the performance of LittleBites on our bulk data, we compared it to two recently described tools designed to extract cell-type specific expression from bulk RNA-Seq data sets. bMIND uses a Bayesian approach to model cell type-specific expression profiles with a pre-calculated proportion matrix, leveraging averaged single cell profiles for each cell type as an informative prior for a gaussian distribution, estimating parameters using a Markov Chain Monte Carlo (MCMC)(*30*). ENIGMA uses a regularized matrix completion approach (using either an L2 regularization or a Trace Norm regularization to estimate a low matrix rank constraint) to estimate expression profiles for each sample(*21*).

Two types of ground truth genes were used for assessment. 1) We used a set of 160 ground truth genes with individual cell-type resolution in the nervous system so that we could also assess how well each technique can distinguish signals expressed in some, but not all, neurons (see Methods; Table S3). 2) We used negative ground truth genes that are expressed exclusively outside of the nervous system (‘non-neuronal’), paired with positive ground truth genes that are expressed in all neuron types (‘ubiquitous’), to assess each method’s ability to accurately remove non-neuronal counts without removing genes expressed in all neuron types (see Methods; Table S4 and S5).

LittleBites was run with a testing subset of ubiquitous and non-neuronal genes as a ground truth comparison in the algorithm, and all assessments between methods were performed using a reserved set of testing genes.

We assessed the performance of LittleBites, ENIGMA, and bMIND against these ground truth genes using the AUROC for the average resulting bulk profiles as the primary metric (Fig. 2B,C; S2A,B). We found that LittleBites improved the ubiquitous and non-neuronal AUROC by 8.7%, ENIGMA improved the neuronal AUROC by 7.8%, and bMIND improved the AUROC by 3.6% (P < 0.0001). Further, LittleBites improved the neuronal AUROC by 7.4%, while ENIGMA and bMIND had a lesser effect.

Comparing results on all individual RNA samples showed consistent improvements across samples from LittleBites (Fig. S2C,D). After LittleBites cleanup, all samples have higher AUROC scores for ubiquitous and non-neuronal testing genes, and all but one sample have higher AUROC scores for the neuronal testing genes compared to unaltered bulk samples. ENIGMA cleanup improves ubiquitous and non-neuronal AUROC for all samples but has little effect on neuronal AUROC for each sample. bMIND improves ubiquitous and non-neuronal AUROC for most samples, but some samples show little effect, and bMIND has a mixed effect on neuronal AUROC, improving some samples while reducing the score for others. We conclude that, for cases where robust ground truth data are available, LittleBites improves the accuracy of cell type expression data in bulk RNA samples.

### A model for the relationship between gene counts and proportions in pseudobulk single-cell RNAseq data

To integrate our previous scRNAseq data(*11*) with our bulk data, we first considered how to analyze the single cell data in each cluster. Gene expression in single cell data can be measured in two ways: as *counts,* averaging or aggregating across all cells in each cluster, or as *proportions,* assessing the proportion of cells in the cluster that detect the gene (Fig. 3A)(*31–33*). Comparing these two measurement systems against a set of ground truth genes in the *C. elegans* nervous system, we found that using the proportion yields a more accurate measurement of gene expression than the alternative of relying on counts (Fig. 3B)(*11*).

**Figure 3:**
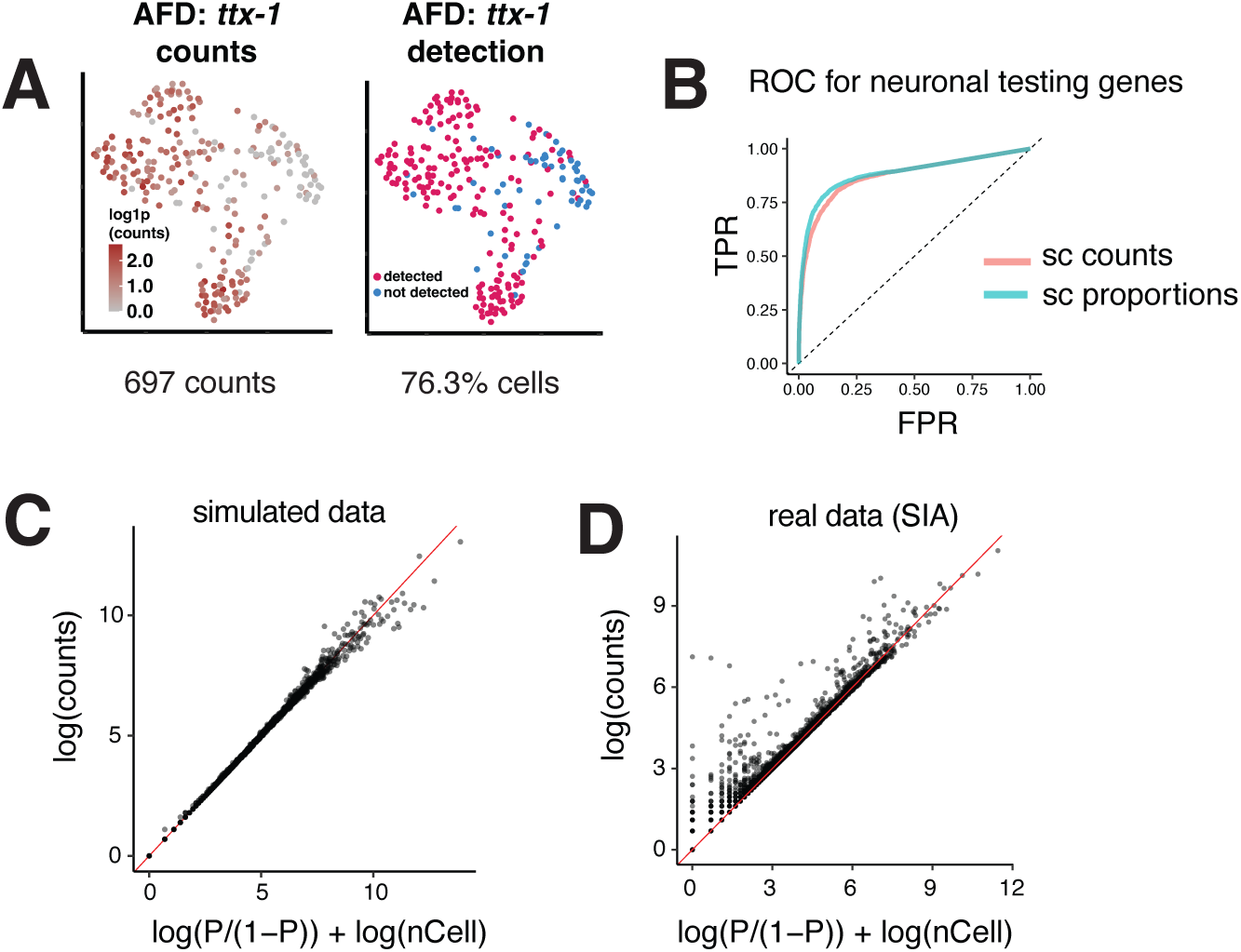
A model for the relationship between cell proportions and gene counts in sc-RNAseq data. A) Aggregate count and proportion measures of scRNA-seq data. UMAP representations of AFD neurons captured in the scRNA-seq dataset(*11*). A) Left, log scale representation of *ttx-1* counts in all AFD neurons. Right, binarized representation of *ttx-1* expression in all AFD neurons. B) ROC for *C. elegans* neuronal ground-truth genes in scRNA-seq counts and proportions. The teal line indicates the proportion of non-zero cells, and the red line indicates the normalized counts. C) Scatter plot of fitted log-counts for simulated data, with log-counts on the Y-axis, and the sum of the log likelihood ratio of the proportion of non-zero cells and the log of the number of cells in the cluster on the X-axis. Red line indicates Y = X. D) Scatter plot of fitted log-counts for one SIA neuron replicate from the *C. elegans* neuronal scRNA-seq dataset(*11*), with log-counts on the Y-axis, and the sum of the log likelihood ratio of the proportion of non-zero cells and the log of the number of cells in the cluster on the X-axis. Red line indicates Y = X.

To exploit the improved accuracy that proportion measures provide, we aimed to transform these proportion values into units of counts for comparison to our bulk data. Building on previous work(*32*), we simulated a single cell dataset, using a negative binomial distribution to model counts for individual cells in a cluster. We found that this simulated dataset could be well-modeled using the log-likelihood ratio (logit) of the non-zero proportion of cells, scaled by the total number of cells in the dataset (Fig. 3C).

Using our real dataset, we observed a similar fit, with the exception that some genes with low non-zero proportions values showed very high counts (Fig. 3D). These exceptions may be caused by high ambient RNA background in some droplets, misassigned cells in the clustering step, doublets that escaped early filtering steps, or rare bursts of expression for lowly expressed genes. The presence of these outliers suggests why the non-zero proportion value may be a more robust measure for gene expression in single-cell clusters than counts: it may be more resistant to rare occurrences of contaminating information.

We used the logit model to generate a denoised counts estimate for each single cell cluster, which we call the scProp2Count dataset. As a final test, we compared the accuracy of the scProp2Count dataset to a standard approach using counts on pseudobulk data, performing pairwise differential expression analysis using edgeR, and then comparing the differentially expressed genes to ground truth expression patterns. We found that the scProp2Count dataset slightly outperforms standard single-cell counts in detection sensitivity and correlation to ground truth, while matching the single-cell counts in precision (Fig. S3).However, caution should be taken when using this approach in scRNAseq cases where all replicates of a cell type contain few cells. scProp2Count values are limited to the space of possible proportion values, and so replicates with low numbers of cells will have fewer potential expression “levels” which may break some model assumptions in downstream applications (see Methods).

### Integrating bulk and single-cell data improves gene detection accuracy

To integrate the bulk and single-cell data sets, for each neuron type in the bulk data set (Table 1) we computed the arithmetic mean of normalized gene counts across all biological replicates (‘Bulk’ dataset). Next, we integrated the Bulk data set with the scProps2Counts dataset by taking the geometric mean for each gene. The result of this analysis was three data sets, each containing normalized gene count data for *C. elegans* neuron classes: Bulk, scProps2Counts, and Integrated. Finally, for each of these three data sets we thresholded the gene counts to call every gene as either expressed or not expressed in each cell type. For this purpose, a single threshold was used for each data set, consisting of the average gene count value for all cells and all genes in each data set. Genes with count values above this threshold were defined as expressed.

To assess whether integration improved the accuracy of expression, we compared expression calls to ground-truth data for both individual neuron types and non-neuronal cell types (see Methods; Table S3,4,5)(*8*). We found that the Integrated dataset generally outperforms the LittleBites-processed Bulk and scProps2Counts scRNA datasets (Fig. 4). Like the LittleBites-processed Bulk dataset, the Integrated dataset approaches a True Positive Rate (TPR) of 100% at relatively low precision (or high FPR), whereas the scProp2Count scRNA dataset has a maximum TPR of 91.9%. The Integrated dataset improves the overall AUROC value by 3.9% compared to the scProp2Count scRNA data, and 2.2% compared to the LittleBites-processed Bulk data (P < 0.0001) (Fig. 4A,B). Like the scProp2Count scRNA dataset, the Integrated dataset has a relatively low False Positive Rate (FPR) for neuronal ground truth genes (Fig. 4C). At a false discovery rate (FDR) of 5%, the Integrated dataset shows substantially better sensitivity than the LittleBites-cleaned Bulk and scProps2Counts scRNA datasets (Fig. 4D). At the level of individual ground truth homeobox and neurotransmitter genes expressed in neurons(*34, 35*), specific and broad improvements can be seen across the Integrated dataset, compared to all other dataset versions (Fig. S4A,B).

**Figure 4:**
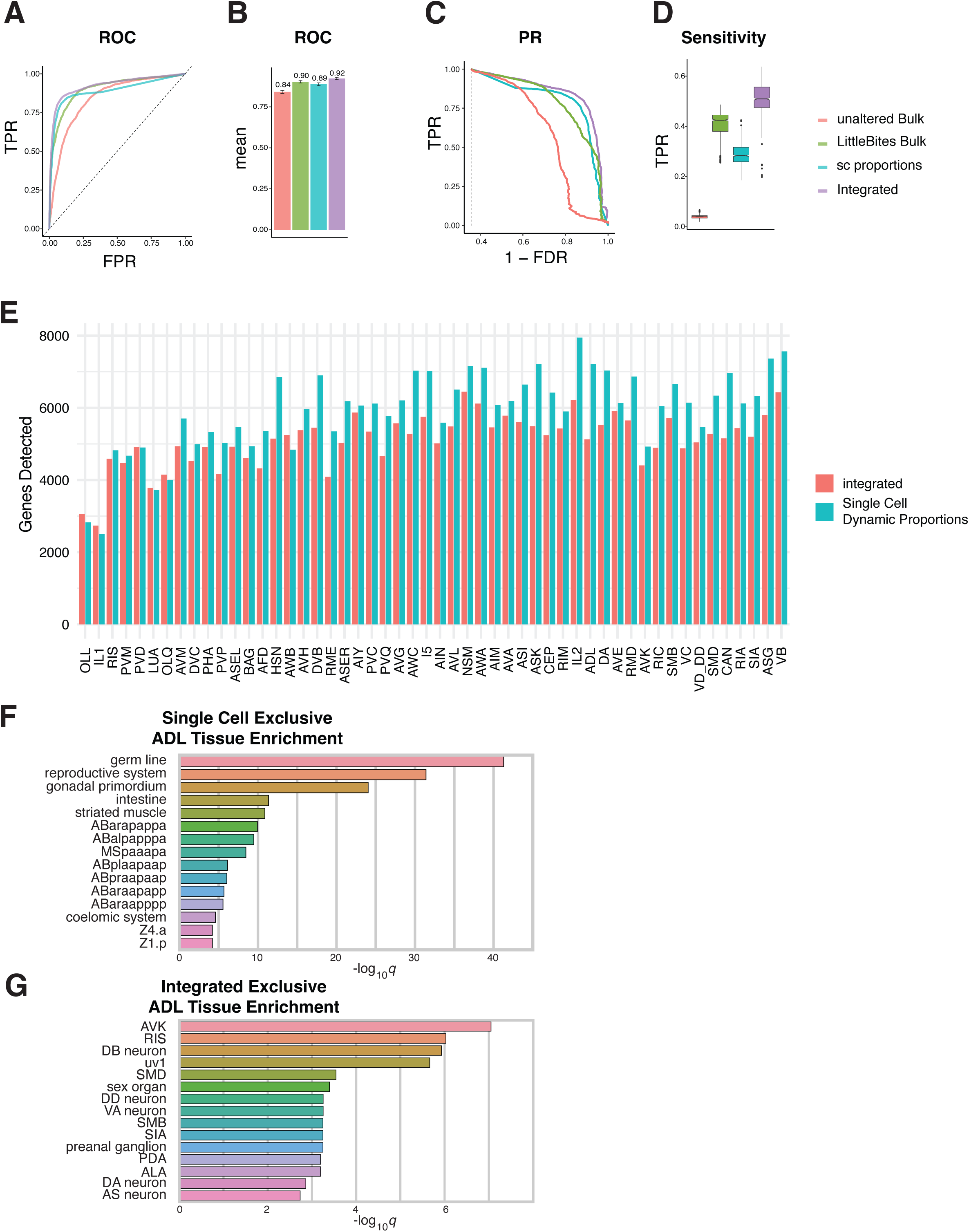
Integrating bulk RNA-seq and scRNA-seq data sets improves gene detection accuracy. A) Receiver Operator Characteristic (ROC) curve for bulk, single-cell, and integrated datasets compared to neuronal ground-truth genes. The x-axis shows the False Positive Rate (FPR), and the y-axis shows the true positive rate (TPR). B) Area under the receiver operator characteristic (AUROC). Error bars represent bootstrap 95% confidence intervals of the AUROC value. Pairwise statistical testing was performed using the deLong method for AUROC scores. *** = p < 0.0001. C) Precision-Recall (PR) curve for bulk, single-cell, and integrated datasets compared to neuronal ground-truth genes. The x-axis shows the Precision (1 – False Discovery Rate/FDR), and the y-axis shows the TPR (Recall). D) Boxplot of bootstrapped sensitivity scores for each dataset at a 5% FDR cutoff. E) Bar chart of the number of genes detected at the medium threshold (FDR = 0.14) for the integrated dataset (red) and the dynamic proportions single cell dataset (blue). Bars are ordered by the number of cells in the corresponding single cell cluster from lowest to highest. F,G) Bar plots of significant terms in Tissue enrichment analyses for genes detected only in the single-cell and integrated datasets in ADL neurons. Dynamic thresholding was used for single cell data, and medium thresholds were used to binarize both datasets.

In addition to using ground truth genes to assess the accuracy of the Integrated dataset, we also evaluated gene expression differences with the original sc-RNAseq dataset(*11*). We thresholded (see Methods) both datasets and determined that the integration strategy results in fewer expressed genes—for some neuron types, over 1,000 fewer genes (Fig. 4E). To further investigate the difference between the Integrated dataset and the original sc-RNAseq dataset, we examined the ‘unique’ genes for each neuron type: i.e., genes called expressed in that neuron type in one dataset but not the other.

We analyzed these ‘unique’ genes using Tissue Enrichment Analysis and Gene Ontology (GO) for two representative neuron types, ADL and VB, both of which are well-represented in the sc-RNAseq dataset(*11*). Importantly, Tissue Enrichment Analysis maps genes to specific *C. elegans* cell types. We found that overall, contamination by other cell types was much lower in the Integrated dataset. Further, the ‘unique’ genes in the sc-RNAseq dataset were more enriched in non-neuronal cell types, such as germline, and for non-neuronal GO terms. By contrast, the ‘unique’ genes in the Integrated dataset were more enriched for other neuronal cell types and neuronal functions (Fig. 4F,G, S4C). Thus, integration appears to successfully remove most contaminating transcripts.

Together, these results show that geometric mean integration of bulk RNA-seq and scRNA-seq datasets combines the strengths of both approaches, providing high sensitivity (bulk RNA-Seq) and high specificity (scRNA-Seq) across a wide range of thresholds. For most purposes, we propose that the Integrated dataset will give the best results. However, for certain applications (such as genes not detected in scRNA-seq data, or non-polyadenylated transcripts, see below) other approaches are needed. To make the Integrated data easily accessible to the community, we provide tables of gene expression for each neuron type based on the Integrated data set, both the values for all genes (Table S6) and values binarized at four thresholds (Tables S7-10).

### Bulk sequencing broadens expression patterns of neuronal genes, particularly for neurons with minimal single-cell data

Our previous scRNA-seq analysis of the *C. elegans* neuronal transcriptome generated a high-density map of protein coding gene expression for a total of 128 transcriptionally distinct neuron types(*11*). However, even this map contains some false negatives— ground truth genes that are known to be expressed in the neuron type but are not detected in the scRNA-seq data. Two factors that contribute to these dropouts are low gene expression and small cluster size, as clusters with few neurons tend to detect fewer genes(*8, 36*). However, our bulk RNA-seq data contains a minimum of 701 cells per bulk sample (Table S1), sequenced to high depth, suggesting that low-expressed genes might be represented in bulk data.

We compared protein coding genes between neuron types in bulk and single-cell data and found a mean Spearman coefficient of 0.612 ± 0.027, with a sharp drop off in the Spearman coefficient for neuron types with the smallest single cell clusters (Fig. 5A). (This analysis used all protein-coding genes detected in a minimum of 3 cells in the single cell dataset.) This result matches previous analysis of the scRNA-seq data, which showed that gene detection is reduced for clusters with < 500 cells(*8*). Together these results suggest that our bulk data may contain gene expression information that is missing from scRNA-seq clusters that contain few cells. If so, this information might be captured in our Integrated dataset.

**Figure 5:**
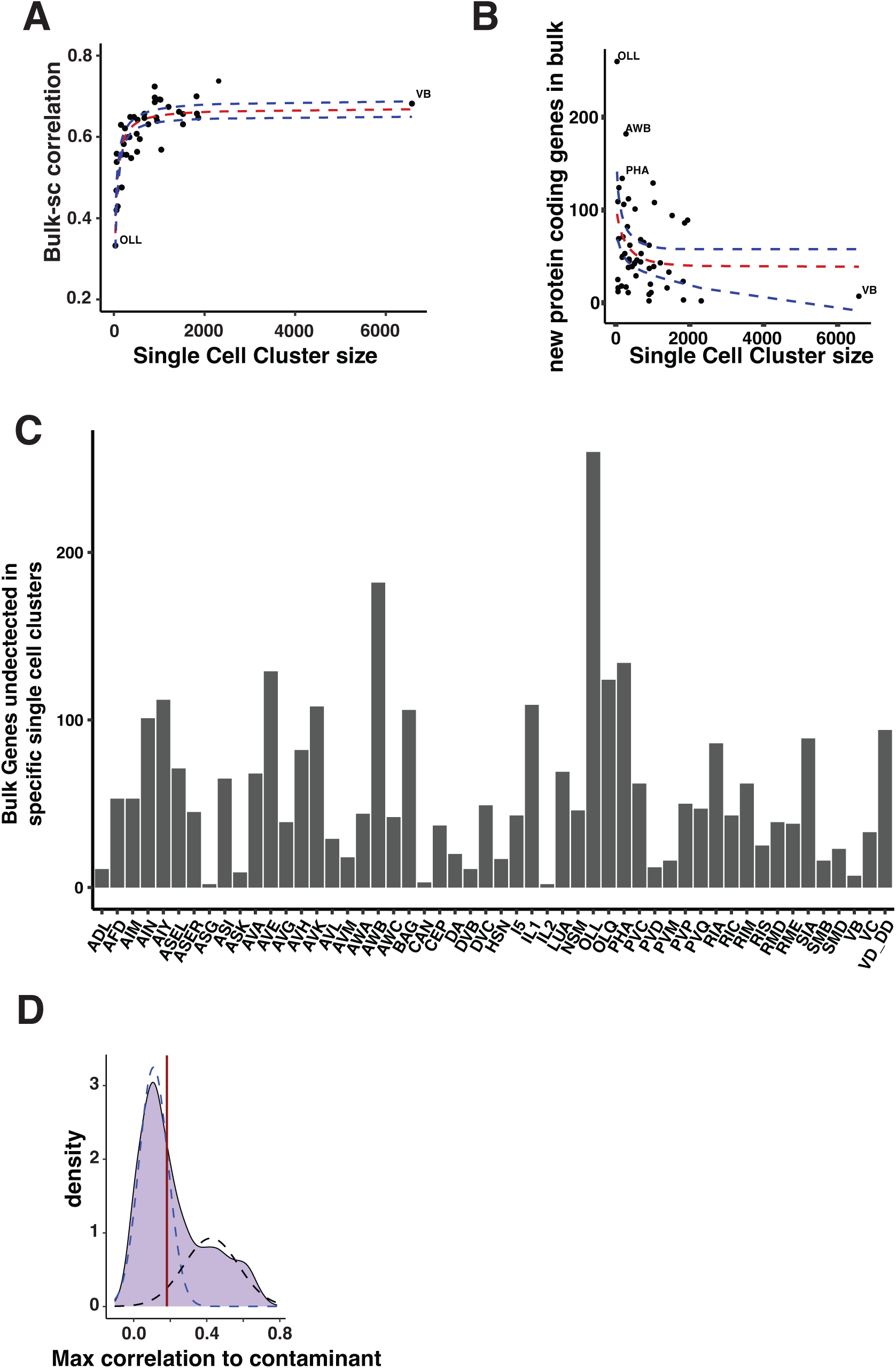
Bulk RNA-seq samples detect protein coding genes that are not detected in scRNA-seq clusters. A) Scatter plot showing the relationship between the size of a scRNA-seq cluster (i.e., the number of cells in the cluster) and the Spearman correlation between the average bulk RNA-seq profile and the average scRNA-seq for all protein coding genes. Each dot represents one cell type. Red dashed line shows a Michaelis-Menten fit (see Methods), gmax = 0.675, beta = 29.507. Blue dashed lines show the 97.5% confidence interval of the fit. B) Scatter plot showing the relationship between the size of a scRNA-seq cluster and the number of additional protein coding genes detected in the integrated dataset per cell type. Each dot represents one cell type. Red dashed line shows an exponential decay fit (see Methods), M = 105.26, m = 44.44, alpha = 238.38. Blue dashed lines show the 97.5% confidence interval of the fit. C) Bar plot showing the number of protein coding genes detected per cell type in the bulk dataset. Genes plotted are: 1) called unexpressed in the corresponding single cell cluster; 2)) are expressed above the medium threshold (FDR = 0.14, 0.237 normalized counts) in the integrated data profile for that cell type. D) Density plot of the gene level correlation to contaminant estimates for all genes that are detected in single cell data. Correlation coefficients were calculated using all unaltered bulk replicates (TMM normalized). Only the highest correlation per gene is used. Blue and black dashed lines represent a Gaussian mixture model, used to threshold against contaminant genes. All protein coding genes with a maximum correlation above 0.175 (red line) were removed from analysis.

We therefore asked whether the Integrated dataset might contain expanded expression information about genes that already have some evidence for expression in scRNAseq data. Because these genes were detected in scRNAseq, they were included in the integrated data analysis. We therefore examined expression of all genes that are detectable in scRNA-seq experiments—in other words, they are called expressed in at least one cell type (by thresholding on the proportion of cells detecting the gene, see Methods). Using a minimum normalized count threshold in the Integrated data to match the FDR of “threshold 2” from the published single cell analysis(*8*), we detected 2 to 260 additional protein coding genes per cell (mean = 58.4, 95% CI = 46.7-72.3). Plotting the number of additional genes against the single cell cluster size reveals that the Integrated dataset detects more additional protein coding genes for cell types with low coverage in the single cell dataset vs cell types with larger numbers of cells in each cluster; the most additional genes were detected in OLL, which has only 34 cells in the single cell dataset (Fig. 5B,C).

We used GO term enrichment to evaluate genes called expressed in the Integrated dataset that were missing in the scRNA-seq data for that neuron type. Most neuron types show enrichment for neuron-associated terms such as neuropeptide signaling, synaptic signaling, cell projection, and gated channel activity. Neuron types with the fewest detected genes in the single cell data also show enrichment for general cell processes such as endosomal functions, chromatin organization, and post-embryonic development (Fig. S5A-E). Thus, the Integrated dataset extends our understanding of neuronal gene expression patterns, with the greatest improvement biased towards clusters with low coverage in the scRNA-seq dataset.

### Bulk sequencing enhances overall detection of neuronal genes

Next, we tested whether the bulk data might yield expression information about genes that were completely undetected in the scRNA-seq dataset. However, since these genes are not included in the scRNA-seq data, the strategies described above— LittleBites and integration, which rely on scRNA-seq data—cannot be used to address the problem of false positives in the bulk dataset derived from non-neuronal tissue contamination. Previous studies have shown that correlations between gene expression and tissue level proportion estimates can be used to deconvolve the profiles of multiple tissues from one mixed bulk profile(*30*). We utilized a similar approach to reduce contamination in our data. First, we estimated contamination in each bulk sample using a non-negative least squares regression (NNLS). We used 100 subsampled datasets to reduce bias against lowly abundant single cell clusters (see Methods). We then calculated per-gene Spearman correlations to each contaminant type (e.g., correlation of *pgl-1* to reproductive cell contamination across all samples). We validated this approach by observing that contaminant correlations for non-neuronal ground-truth genes are higher than the contaminant correlations for all other protein coding genes (Fig. S5F). Using the highest correlation per gene, we modeled this data as a mixture of two Gaussian distributions, one distribution of low contamination correlation scores representing truly expressed neuronal genes, and a second distribution of higher contamination correlation scores representing genes likely detected due to contamination from non-neuronal tissues. (Fig. 5D). Setting a threshold which removes all genes with a contaminant correlation higher than 0.175 excludes 95% of the predicted contaminant distribution profile.

We identified genes whose expression was not detected in sc-RNAseq data in two ways. First, we selected the 873 protein coding genes that were excluded from analysis in the scRNA-seq dataset because they were detected in fewer than 3 of the 100,955 cells sequenced(*11*). Second, we applied a conservative threshold to identify 3,567 protein coding genes that were included in the scRNA-seq analysis but that are not called ‘expressed’ in any cell types, including non-neuronal tissues (see Methods). We combined these gene sets to generate a list of 4,440 ‘unexpressed’ genes that were not detected in the single cell analysis (Table S11).

To examine expression of these unexpressed genes in the bulk data, we first ‘decontaminated’ the bulk data by removing genes with strong correlations to any contaminants as described above. We used the non-neuronal ground-truth genes to set a minimum normalized counts threshold for calling expression, which was set to a non-neuronal FPR of 0%. Using this threshold on the remaining decontaminated unexpressed genes, we detected between 3 and 161 new protein coding genes per cell type. Using ADL as an example, we performed Tissue Enrichment Analysis on the 161 new genes(*37*). The most enriched term is “ADL genes”, as expected, followed by the “amphid sensillum” and “lateral ganglion”, structures that include the ADL neuron (Fig. S5G)(*38*). Thus, our analysis of bulk RNA-Seq data complements our previous scRNA-Seq study(*11*) by identifying likely instances of gene expression that were not detected by scRNA-seq.

### Bulk RNA-seq reveals both broadly expressed and neuron-specific noncoding RNAs

A significant benefit of our bulk RNA-seq approach is its sensitivity to non-poly-adenylated transcripts, which include many species of non-coding RNA(*9*). However, similar to coding genes not found in the scRNA-seq dataset (see above), the scRNA-seq dataset is largely uninformative about these transcripts. Further, we do not have a ground-truth data set of non-coding genes to evaluate accuracy of expression calling for these transcripts. Finally, most non-coding RNAs are expressed at lower levels than protein coding genes (Fig. S6A). We therefore applied a generous, uniform threshold for calling gene expression, consisting of > 5 normalized counts in 2/3 or more of the samples within a cell type. As we did for our analysis of coding genes not found in scRNA-seq, we used gene level correlation to contamination estimates as a procedure to eliminate non-coding genes that were likely detected due to contamination from other tissues in the bulk samples. First, we estimated contamination for each sample using a bootstrapped NNLS regression (Fig. S6B; see Methods), and then calculated per-gene Spearman correlations to each contaminant type. We applied a threshold on the gene level correlation to contamination estimates for each sample by fitting a Gaussian mixture model to the maximum correlation score for each gene. We selected a cutoff of 0.23, which excludes 98% of the estimated contamination distribution (Fig. S6B). With these thresholds, an average of 512 noncoding RNAs were identified as “expressed” per cell type (95 CI ± 40.6) (Table S12). By RNA type, we detected 16.2 ± 1.2 lincRNAs, 39.0 ± 5.3 pseudogenes, 54.8 ± 9.7 tRNAs, 41.8 ± 2.0 snRNAs, 131.5 ± 1.8 snoRNAs, and 228.9 ± 29.0 uncategorized ncRNAs per cell type (Table S13).

Next, we sought to identify noncoding RNAs with broad expression across multiple neuron types. We defined broad expression as genes expressed in > 80% of neuron classes in our dataset (Fig. S6C). This approach defined 247 non-coding genes as broadly expressed, or “pan-neuronal”. These pan-neuronal noncoding RNAs include 108 (43.8%) snoRNAs and 33 (13.4%) snRNAs, both tenfold greater than the expected proportion assuming a random distribution (Fisher’s exact test, P-value < 0.01) (Fig. 6A). By contrast, pseudogenes and otherwise uncategorized ncRNAs were significantly depleted from the list of pan-neuronal noncoding RNAs (P-value < 0.001). These results indicate that snoRNAs and snRNAs are widely expressed, which matches studies showing broad expression of many snoRNAs and snRNAs in other systems(*39, 40*), and is consistent with their key roles in rRNA processing and splicing(*41–43*).

**Figure 6:**
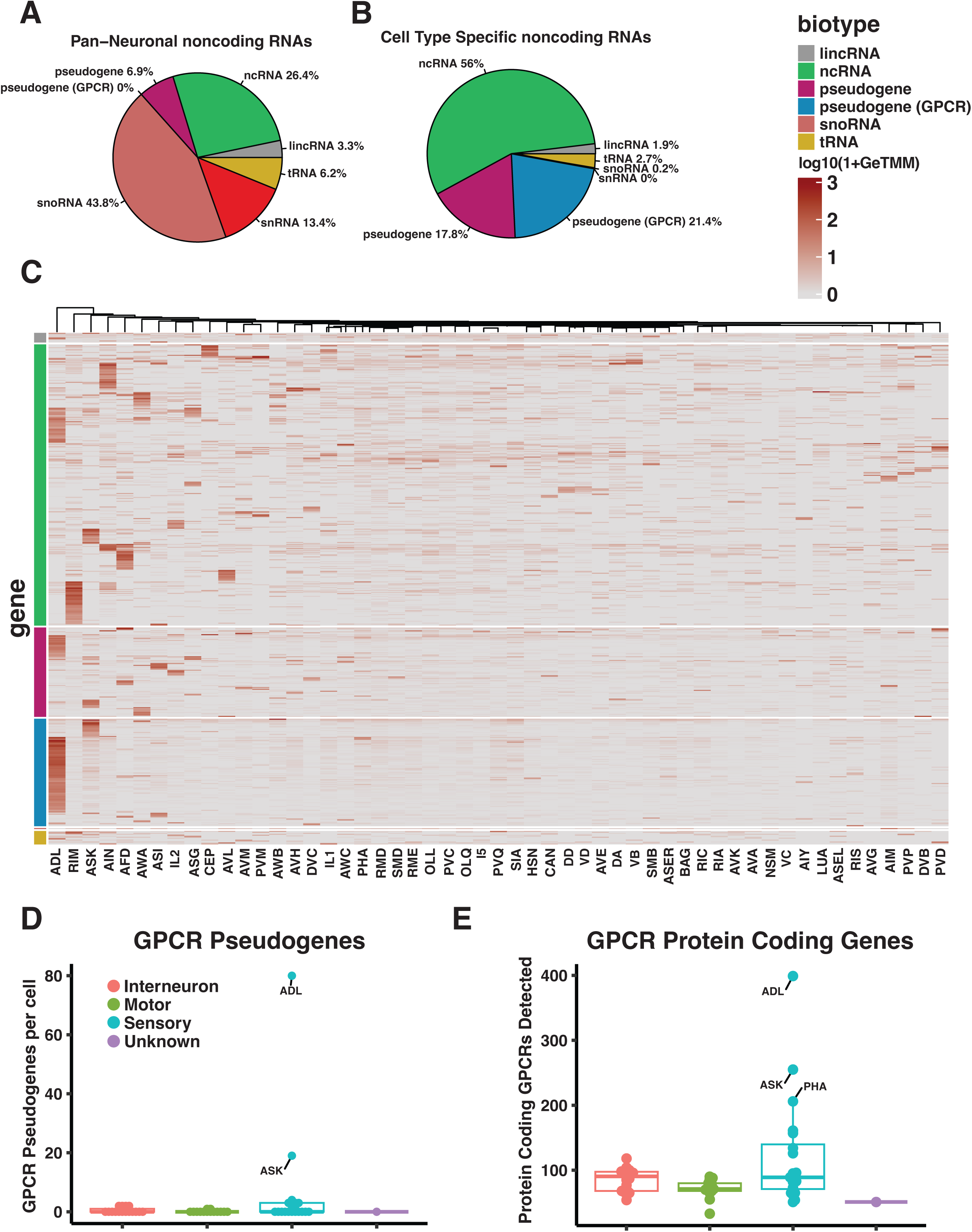
Bulk analysis reveals noncoding RNA expression patterns. A) Proportions of classes of pan-neuronal noncoding RNAs. B) Proportions of classes of cell type specific noncoding RNAs C) Heatmap of average normalized counts per cell type for ncRNA genes considered cell type specific, rows are grouped by RNA class. Note that genes within individual neuron types are clustered by expression patterns, not similar gene function. D) Number of GPCR pseudogenes with specific expression per neuron type, grouped by neuron function. E) Number of GPCR genes per neuron type, grouped by neuron function.

We also sought to identify neuron-type-specific noncoding RNAs. We calculated tissue specificity scores for each noncoding RNA called expressed in at least one neuron type using the Preferential Expression Measure (PEM) score(*23, 44*). We called these genes neuron-type specific according to three criteria: (1) Called expressed in ≥ one neuron type (see above); (2) PEM score > 0.65; (3) > 2 normalized counts in a maximum of 10/53 neuron types. Using these thresholds, we identified 523 cell-type-specific noncoding RNAs (Fig. 6B). By RNA type, 293 (56.0%) of cell type-specific noncoding RNA genes are uncategorized ncRNAs, 93 (17.8%) are general pseudogenes 112 (21.4%) are GPCR pseudogenes, 14 (2.7%) are tRNAs, 10 (1.9%) are lincRNAs, 1 (0.2%) is a snoRNA, and none are snRNAs. We observed significant enrichment of pseudogenes, and a small but significant depletion of ncRNAs, snoRNAs, and tRNAs (P-value < 0.01). Clustering by genes and cell type modalities revealed clear enrichment for specific noncoding RNAs in individual neuron types (Fig. 6C), although the overall proportions of noncoding RNA types were similar between individual neuron classes (Fig. S6D). The number of specific noncoding RNAs per cell type ranged from 0 (PVC) to 139 (ADL), with a mean of 10.2 (± 5.8).

Interestingly, we observed that in ADL—the neuron that expresses the highest number of specific ncRNAs—many of these specific ncRNAs are GPCR pseudogenes (Fig. 6C,D, Table S14). ADL also expresses the highest number of bona fide GPCRs among neurons in our dataset (Fig. 6E, Table S15). Looking across neuron types, we observe a rough correspondence between expression of specific GPCR pseudogenes and expression of bona fide GPCRs (Fig. 6D,E). This analysis suggests that pseudogenes retain their regulatory information, despite losing their coding potential. Overall, our data on expression of non-coding genes reveal a wide diversity of noncoding RNA expression across the nervous system and open the door to in depth studies of noncoding RNA contributions to individual neuron function.

## Discussion

In this work, we present bulk RNA-seq data for 53 neuron classes in the *C. elegans* nervous system. We describe new methods for integrating these bulk RNA-seq data with previously obtained single-cell RNA-seq data(*11*). We find that this approach improves gene detection accuracy in comparison to each individual data set. We use the bulk data sets to detect two important classes of genes that were underrepresented in the previous single-cell atlas: (1) lowly expressed genes, and (2) non-poly-adenylated transcripts, chiefly non-coding RNAs. Overall, our results provide a rich resource for studies of gene expression in the *C. elegans* nervous system.

The rapid growth of scRNA-seq and bulk RNA-seq datasets create opportunities to combine these two data types to address diverse goals. For example, recent studies have combined data from bulk RNA sequencing and scRNA-seq to address the problem of deconvolution. The goal of deconvolution is to infer cell-type expression profiles from tissue level bulk samples, using scRNA-seq references as a guide(*30, 45–47*). In this study we combine these two types of data, both collected from the *C. elegans* nervous system, for a different purpose: to improve the accuracy of cell and tissue-specific transcriptional profiles. We developed two approaches: a subtraction approach, LittleBites, that uses information from scRNA-seq data, combined with ground truth gene expression data, to identify and remove contaminating counts from bulk RNAseq data; and an integration approach that directly combines information from bulk and single-cell approaches to improve sensitivity and specificity. Our computational approaches may be useful in other cases where scRNA-seq and bulk RNA-seq datasets are available to improve information on gene expression.

In addition to enhancing the accuracy of gene expression, the integrated bulk RNA-seq dataset detects lowly expressed protein coding genes that were not detected by scRNA-seq. Further, because our library construction methods were designed to capture non-polyadenylated transcripts, our bulk RNA-seq data set detects noncoding RNAs that were not revealed by previous scRNA-seq results(*8, 9*). Some of these noncoding RNAs are broadly expressed in the nervous system, suggestive of shared functions across different types of neurons. Other non-coding RNAs are expressed in a limited number of neuron types, suggesting neuron type-specific functions. In addition, the bulk RNA-seq dataset contains transcript information across the gene body, which can yield information about mRNA splicing that is not found in the scRNA-seq dataset(*26, 27*).

Overall, our approach achieves a comprehensive representation of all classes of transcripts expressed in individual neuron types. These data can now drive analysis of mechanisms that control gene expression across the genome in individual neuron types, and support identification of differentially expressed genes that define neuron-type specific differences in morphology and function. Public access to these data (described below) will enable further analysis into the regulation and function of differential gene expression in *C. elegans* neurons.

## Data Availability

Bulk raw data and single cell raw data are available at Gene Expression Omnibus (GEO, https://www.ncbi.nlm.nih.gov/geo). Bulk data identifier: GSE229078. Single cell data identifier: GSE136049. Counts data and additional supporting files can be downloaded from the CeNGEN website (https://www.cengen.org). Code is available at GitHub (https://www.github.com/cengenproject).

## Figures and Tables

**Table 1:**

All strains used for bulk RNA-seq experiments, including strain names, allele information, and transgenic fluorescent labels for sorting individual neuron types.

**Supplementary Table 1:**

Replicate metadata for bulk RNA-seq experiments, with replicate names, strain names, and the number of cells collected.

**Supplementary Table 2:**

List of genes that were removed from analysis because they overlap with promoter regions used in the construction of transgenic markers for FACS sorting.

**Supplementary Table 3:**

Ground Truth expression matrix for 160 genes in the *C. elegans* nervous system using fosmid and CRISPR/Cas reporter lines (see Methods).

**Supplementary Table 4:**

List of 445 genes that are expressed exclusively outside the *C. elegans* nervous system (see Methods).

**Supplementary Table 5:**

List of 936 ubiquitous genes (see Methods).

**Supplementary Table 6:**

Integrated gene expression matrix.

**Supplementary Tables 7-10:**

Thresholded gene-by-cell expression matrices using the Integrated data set, computed at four thresholds, values below the threshold are set to 0.

**Supplementary Table 11:**

List of genes called unexpressed in all single cell clusters (see Methods).

**Supplementary Table 12:**

List of ncRNA genes expressed in individual neuron types.

**Supplementary Table 13:**

List of ncRNA gene types expressed in individual neuron types.

**Supplementary Table 14:**

List of GPCR pseudogenes included in this analysis.

**Supplementary Table 15:**

List of protein coding GPCR genes included in this analysis.

---

**Supplementary Figure 1.**
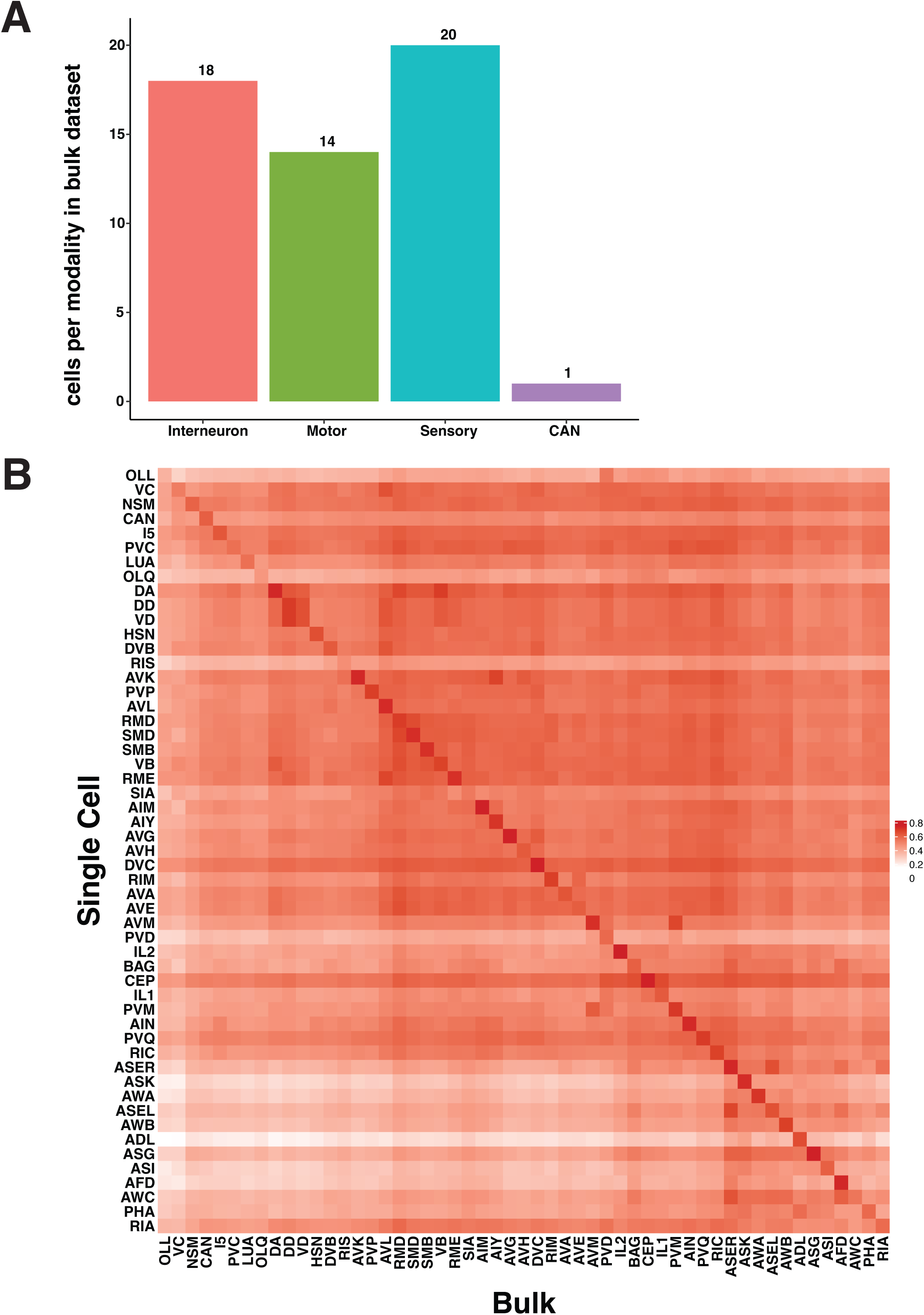
A) Number of cell types sequenced per functional modality. B) Heatmap of Spearman Correlations between average single cell RNA-seq (row) and Bulk RNA-seq (column) profiles for each neuron type. For each row, correlations were calculated for genes called expressed in that single cell cluster (from single cell thresholding)(*11*). The single cell data did not separate the DD and VD GABA motor neuron types(*11*); these rows are identical.

**Supplementary Figure 2.**
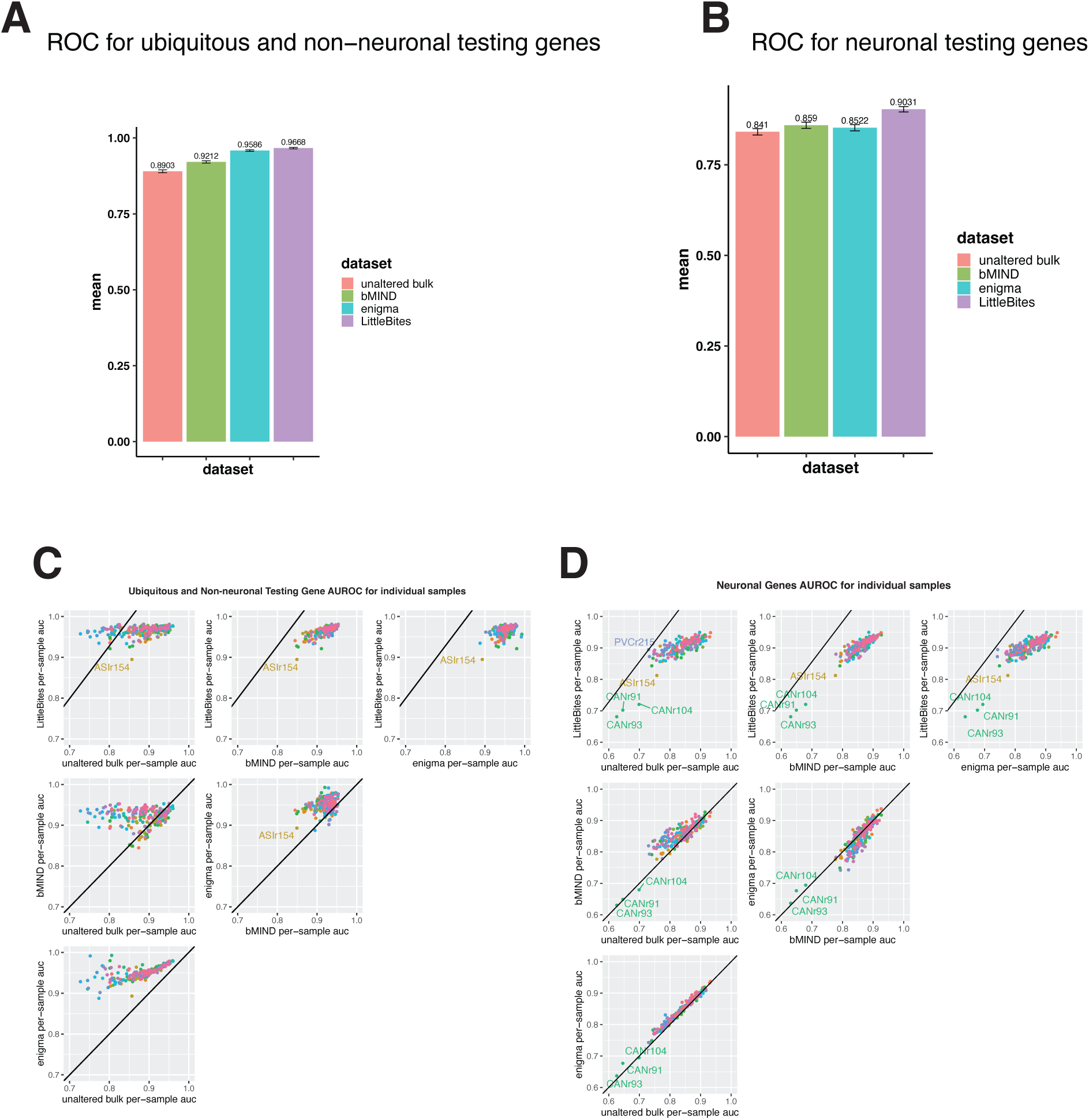
A) Bar graph showing the AUROC values for each dataset assessed against ubiquitous genes and genes exclusively expressed in non-neuronal cells. Error bars represent bootstrap estimated confidence intervals. B) Bar graph showing the AUROC values for each dataset, assessed against 160 genes expressed within the nervous system. Error bars represent bootstrap estimated confidence intervals. C,D) Scatter plots of pairwise comparisons of sample-level AUROC values for unaltered and corrected bulk datasets. Ubiquitous and non-neuronal AUROCs are represented in panel C, and neuronal AUROCs are represented in panel D.

**Supplementary Figure 3.**
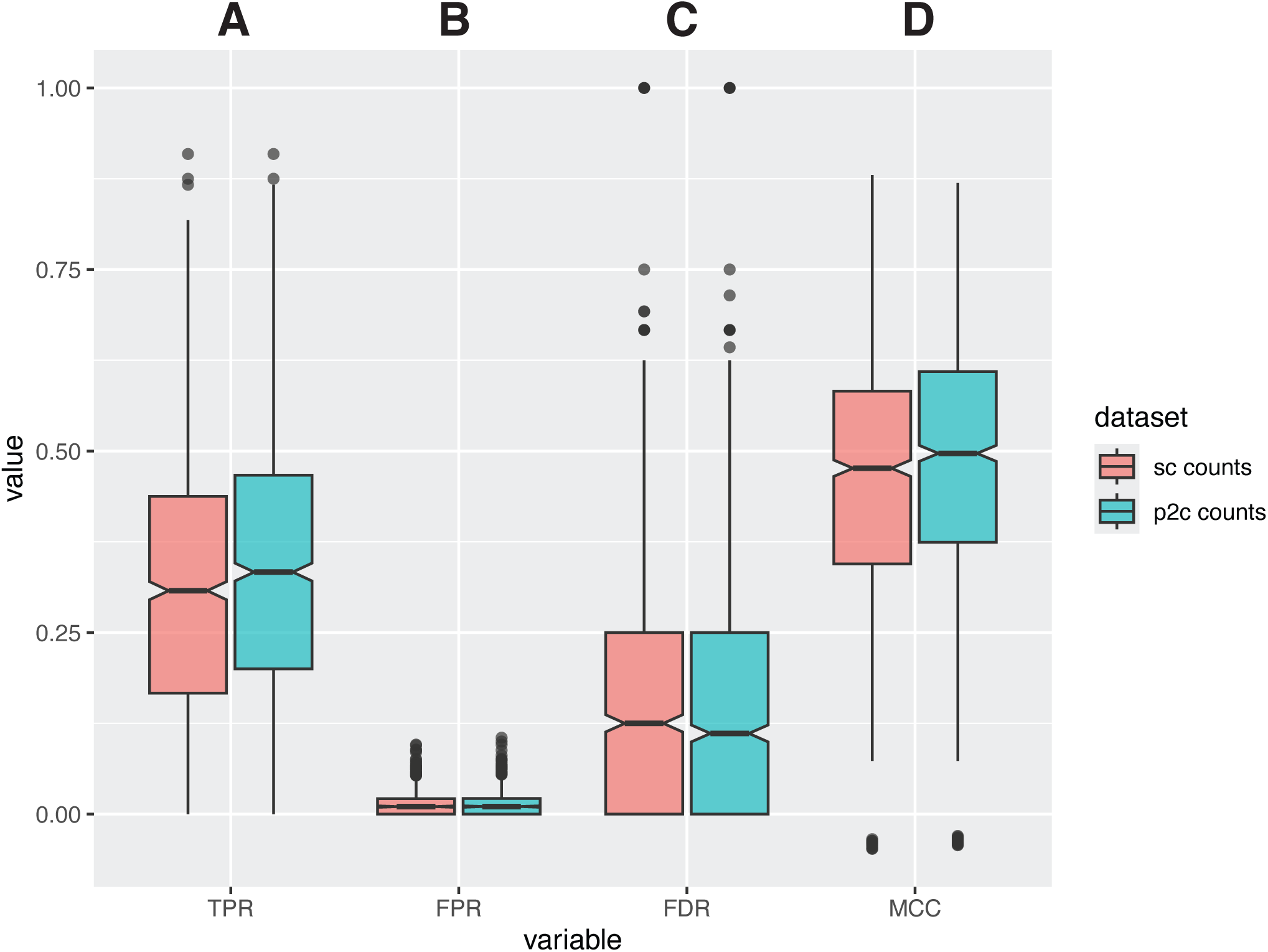
Box plots of edgeR differential expression metrics for single-cell counts and Prop2Count inputs. 1176 neuron-neuron pairs were compared in edgeR, and differentially expressed genes were called as any gene with a p.adj value < 0.05, and a log2 effect size > 2. Metrics were generated by comparing differentially expressed genes with 160 differentially expressed ground truth genes. A) Box plots of TPR values for neuron-neuron edgeR comparisons. B) Box plots of FPR values for neuron-neuron edgeR comparisons. C) Box plots of FDR values for neuron-neuron edgeR comparisons. D) Box plots of MCC values for neuron-neuron edgeR comparisons. Red box plots represent single-cell counts edgeR comparisons, and teal box plots represent Prop2Count edgeR comparisons. All comparisons were made with a two-way permutation test (* = p < 0.05, ** = p < 0.01).

**Supplementary Figure 4.**
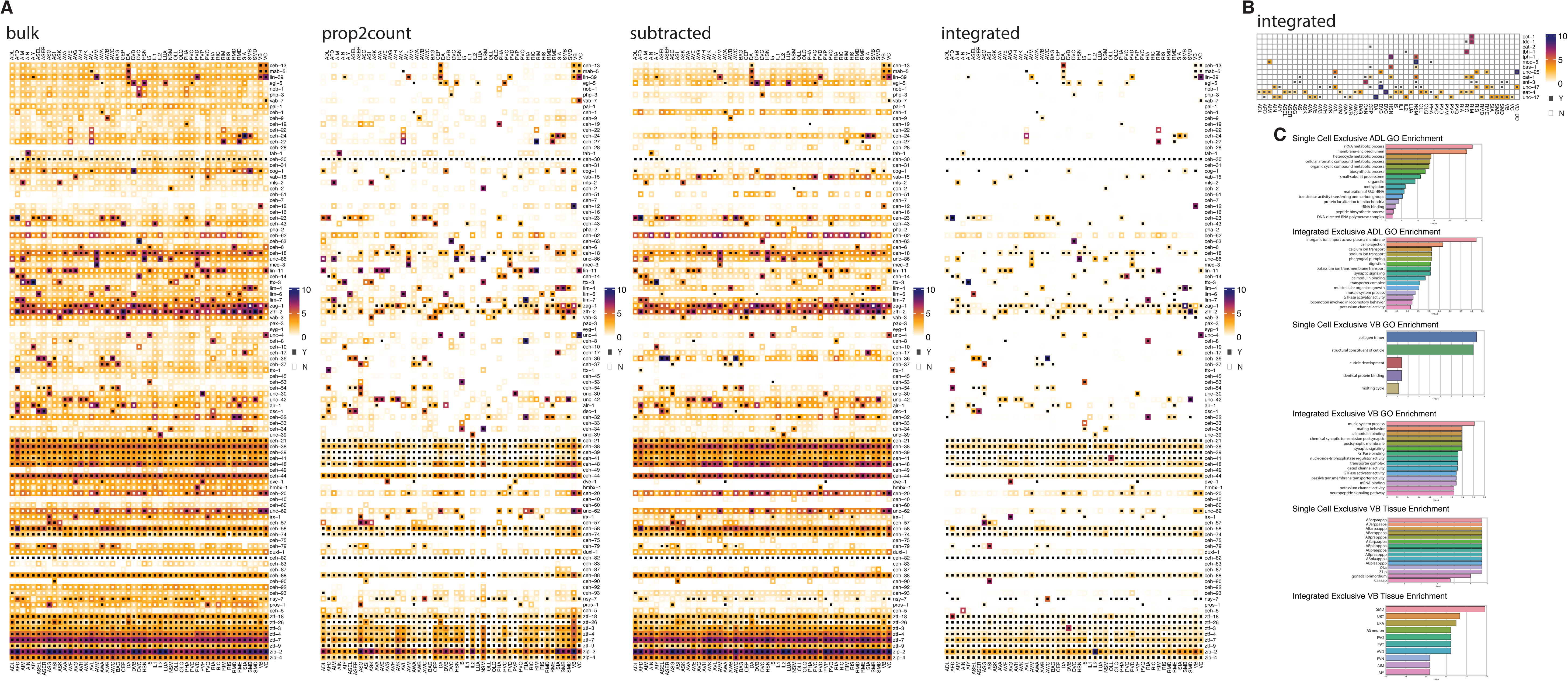
A) Comparison between ‘ground truth’ fluorescent reporters and RNAseq gene expression for all datasets, shown for a set of homeobox genes(*34*). Fluorescent reporter expression is binarized and is shown as small central squares, with black = expressed and white = not expressed. RNA transcripts shown as natural log transformed TPM counts, color-mapped according to the scale. B) Comparison between ‘ground truth’ fluorescent reporters and RNAseq gene expression for the Integrated dataset, shown for a set of genes encoding neurotransmitter machinery(*35*). Fluorescent reporter expression is binarized and is shown as small central squares, with black = expressed and white = not expressed. RNA transcripts shown as natural log transformed TPM counts, color-mapped according to the scale. C) Bar plots of significant terms in GO and Tissue enrichment analyses for genes detected only in the single-cell and integrated datasets in ADL and VB neuron types.

**Supplementary Figure 5.**
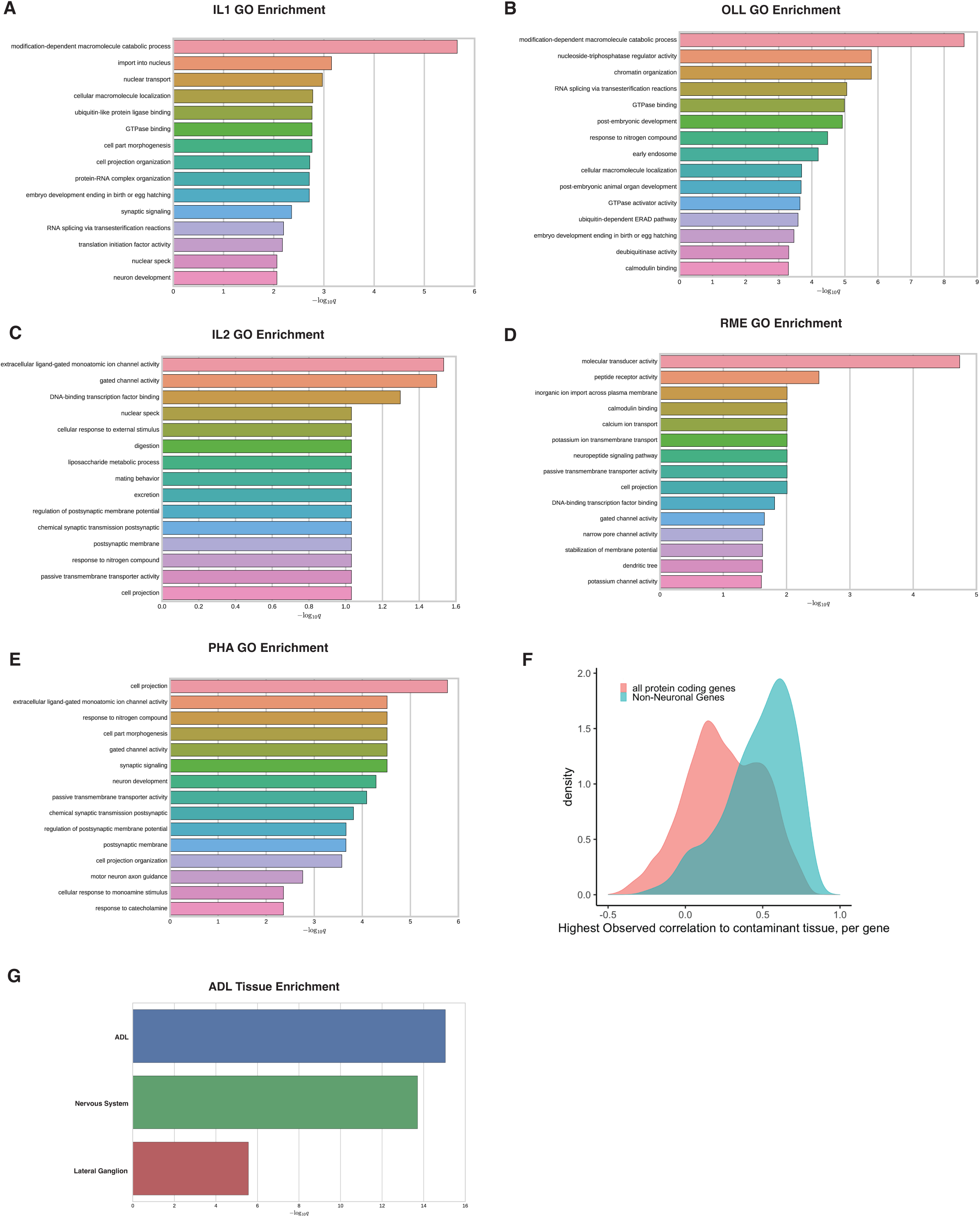
A-E) GO enrichment analysis for protein coding genes detected in selected bulk samples that were not detected in the corresponding scRNA-seq cluster. GO enrichment performed using WormBase. F) Density plot gene level correlations to non-neuronal contamination estimates in bulk samples. Maximum correlation per gene is plotted. Blue density curve represents a set of genes identified as putative non-neuronal markers, and red density curve represents all other protein coding genes, only protein coding genes called expressed in at least one neuronal cell type were plotted. G) Tissue Enrichment Analysis of 161 ‘new’ genes found in ADL.

**Supplementary Figure 6.**
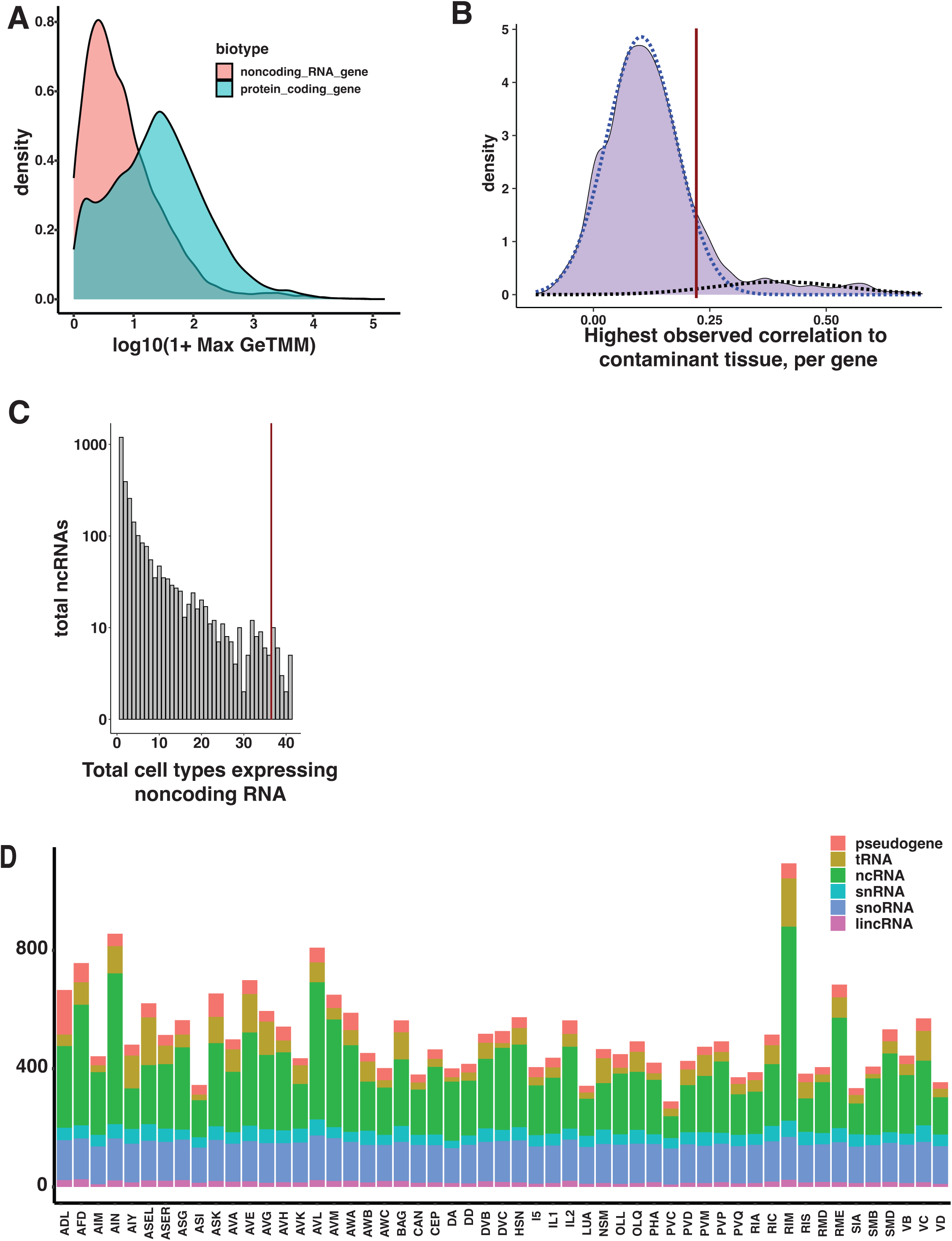
A) Log-scaled expression distribution of protein coding genes (blue) and non-coding genes (pink). B) Density plot showing the distribution of gene level correlation to contaminant estimates (purple); values plotted are the highest correlation per gene. Genes plotted were called expressed in at least one cell type. Blue and black dashed lines represent a gaussian mixture model, used to threshold against contaminant genes. All noncoding genes with a maximum correlation above 0.22 (vertical red line) were removed from analysis. C) Histogram of ncRNA genes binned by number of neuron types in which they are expressed; genes to the right of the red line are expressed in >90% of neuron types in this study. D) Barplot of the number of noncoding RNAs detected in each neuron type, colored by noncoding RNA class.

